# Inhibition of c-Jun in AgRP neurons mediates chronic stress-induced anxiety-like behaviors and colitis susceptibility

**DOI:** 10.1101/2022.05.02.490271

**Authors:** Fuxin Jiao, Xiaoming Hu, Hanrui Yin, Feixiang Yuan, Ziheng Zhou, Wei Wu, Shanghai Chen, Zhanju Liu, Feifan Guo

## Abstract

Psychiatric disorders, such as anxiety, are frequently associated with inflammatory bowel diseases (IBD), however, the neural mechanisms are unknown. Here, we showed that hypothalamic agouti-related protein (AgRP) neuronal activity was suppressed under chronic restraint stress (CRS), a condition known to induce anxiety-like behaviors and increase colitis susceptibility. Consistently, chemogenic activation (inhibition) of AgRP neurons reversed (mimicked) CRS-induced anxiety-like behaviors and colitis susceptibility. Furthermore, CRS inhibited AgRP neuronal activity by suppressing the expression of c-Jun. As expected, overexpression of c-Jun in these neurons protected against the CRS-induced these effects and knockdown of c-Jun in AgRP neurons (*c-Jun*^ΔAgRP^) promoted anxiety-like behaviors and colitis. Moreover, relieving the anxiety with cyamemazine (an anxiolytic drug) alleviated colitis susceptibility in *c-Jun*^ΔAgRP^ mice. Finally, according to a proteomic analysis, the levels of the secreted protein thrombospondin 1 (THBS1) were negatively associated with the increased anxiety-like behaviors and colitis susceptibility, supplementing recombinant THBS1 rescued colitis in *c-Jun*^ΔAgRP^ mice. Taken together, these results reveal a critical role of hypothalamic AgRP neuron-derived c-Jun in orchestrating chronic stress-induced anxiety-like behaviors and colitis susceptibility. These results provide a new perspective for understanding the neuronal mechanisms and potential therapeutic target for the comorbidity of psychiatric disorders, such as anxiety, and IBD.

## Introduction

Psychological disorders, such as anxiety and depression, severely influence human health (Kalin, 2020). What makes it even worse is that they are frequently associated with the occurrence of many other diseases, including inflammatory bowel diseases (IBD) (Kurina et al., 2001; Wang et al., 2020; Xia et al., 2021). Stress is a common cause of psychiatric disorders (de Kloet et al., 2005) and intestinal inflammation (Gao et al., 2018; Qiu et al., 1999). Accumulating lines of evidence has illustrated that brain-gut interactions play an important role in the outcome of psychological disorders and IBD under stress conditions (Bonaz and Bernstein, 2013; Gracie et al., 2019). Although gut signals and microbiota are proposed to play roles in the comorbidity of psychological disorders and intestinal inflammation (Gao et al., 2018), the neural mechanisms behind stress-induced anxiety and colitis susceptibility are unknown.

The hypothalamus is an important neural control center in the regulation of the stress response (Bains et al., 2015), consisting of a series of important nuclei. The hypothalamic arcuate nucleus (ARC) is critical to the regulation of energy metabolism (Sohn et al., 2013) and recently reported to be engaged in emotional regulation (Fang et al., 2021; Qu et al., 2020; Xia et al., 2021). There are two specific populations of neurons in the ARC: neurons co-expressing the orexigenic neuropeptide agouti-related protein (AgRP) and neuropeptide Y, and neurons co-expressing the anorexigenic pro-opiomelanocortin (POMC) precursor and the cocaine- and amphetamine-related transcript, whereas the AGRP/NPY neurons have an inhibitory effect on the POMC/CART neurons (Bell et al., 2005). Unlike the POMC that is widely produced in addition to ARC (Harno et al., 2018), neurons expressing AgRP are localized exclusively in the hypothalamic ARC (Broberger et al., 1998). AgRP neuronal activity is influenced by stress (Fang et al., 2021), and controls diverse physiological processes, including feeding (Bell et al., 2005), pain sensation (Alhadeff et al., 2018) and food-seeking behavior (Dietrich et al., 2015). Studies shows that AgRP neurons are involved in the regulation of peripheral tissue homeostasis (Joly-Amado et al., 2012; Kim et al., 2015), suggesting that it may play an important role in stress-induced anxiety and colitis.

c-Jun is a component of the activator protein-1 transcription factor family, which forms either a homodimer or heterodimer with other members of the family (c-Fos or ATF) and plays a role in the activation of downstream target genes (Wisdom et al., 1999). It is expressed in many tissues, including brain (Sakai et al., 1989). c-Jun is shown to be involved in the regulation of many functions in peripheral tissues (Fuest et al., 2012; Nateri et al., 2005). By contrast, the role of c-Jun in the brain is poorly understood, except for some biological processes such as axonal injury (Raivich et al., 2004) and neurodegeneration (Raivich and Behrens, 2006). Because c-Jun is an immediate-early gene that is dynamically regulated in response to neuronal activity (McNeill and Robinson, 2015), it is commonly used as a marker reflecting neuronal activity (Hoffman et al., 1993). However, it is also induced under stress conditions (Filipovic et al., 2012) and highly expressed in the hypothalamus arcuate nucleus (Herdegen et al., 1995), suggesting that it may additionally be involved in the regulation of some important functions in the arcuate nucleus.

Based on the above knowledge, we hypothesize that c-Jun in AgRP neurons plays an important role in the stress-induced comorbidity of anxiety and IBD. This study investigates such a possibility and explores the likely mechanisms.

## Results

### Chronic Restraint Stress (CRS) Induces Anxiety-Like Behaviors and Increases Susceptibility to Colitis

To induce anxiety and colitis, we employed a CRS mouse model (Supplementary information Fig. S1A) as described previously (Gao et al., 2018; Liu et al., 2020; McGill et al., 2006), which significantly reduced body weight, elevated adrenal gland weight and serum corticosterone levels compared with control treatment (Supplementary information Fig. S1B-D). As shown previously (Liu et al., 2020), CRS induced anxiety-like behaviors, as demonstrated by the significantly reduced time and travel distance in the central region in the open field (OF) test, and the shorter time and fewer entries to the open arms in the elevated plus maze (EPM) test (Supplementary information Fig. S1E and F). In addition, the CRS mice exhibited a greater extent of dextran sodium sulfate (DSS)-induced colitis, as evaluated by the loss of body weight, gross bleeding, and shortening of colon length, as well as histological analysis revealing epithelial damage and lymphocyte infiltration of the distal colon (Supplementary information Fig. S1G-J). Consistently, the mRNA levels of proinflammatory cytokines (*interleukin (IL)* 6 (*Il6), Il1b, Il12*, and *transforming growth factor beta (Tgfb)* (Tian et al., 2019) were significantly increased in the colon tissues of CRS mice (Supplementary information Fig. S1K). These results suggest that CRS could induce anxiety-related behaviors and colitis.

### Activation of AgRP Neurons Reverses CRS-increased Anxiety-like behaviors and Colitis Susceptibility

To investigate the involvement of AgRP neurons in CRS-induced effects, we conducted immunofluorescence (IF) staining for c-Fos, a signal reflecting neuronal activity (Krashes et al., 2011), in the AgRP-Cre-Ai9 mice. IF staining of tdTomato (reflecting AgRP neurons) and c-Fos revealed decreased c-Fos levels in the AgRP neurons of CRS mice (Supplementary information Fig. S2A), suggesting inhibited AgRP neuronal activity. If the inhibited AgRP neurons were important in this case, stimulating AgRP neurons by an excitatory DREADD receptor hM3Dq (Krashes et al., 2013), should reverse the CRS-increased susceptibility to anxiety and colitis. As predicted, stimulation of AgRP neurons (as shown by the increased c-Fos staining, Supplementary information Fig. S2B and C) reversed CRS-induced anxiety-like behaviors with an increase in both center time duration and center distance in the OF test (Fig. 1A), and an increase in time and entries into the open arms in the EPM test (Fig. 1B). Furthermore, the activation of AgRP neurons also reversed chronic stress-increased susceptibility to DSS-induced colitis, as demonstrated by its blocking effects on the CRS-induced loss of body weight, increased bleeding score, shortened colon length, higher histological scores, and increased expression of proinflammatory factors (*Il6, Il1b, Il12*, and *Tgfb)* (Fig. 1C-G, Supplementary information Fig. S2D). Although activation of AgRP neuron has a significant impact on feeding behavior (Krashes et al., 2011), the effect of colitis-related findings was not due to food intake as shown by pair-fed experiments (Supplementary information Fig. S3A-E). These results suggest that activation of AgRP neurons enables to reverse anxiety-related behaviors and colitis induced by CRS.

**Figure 1.**
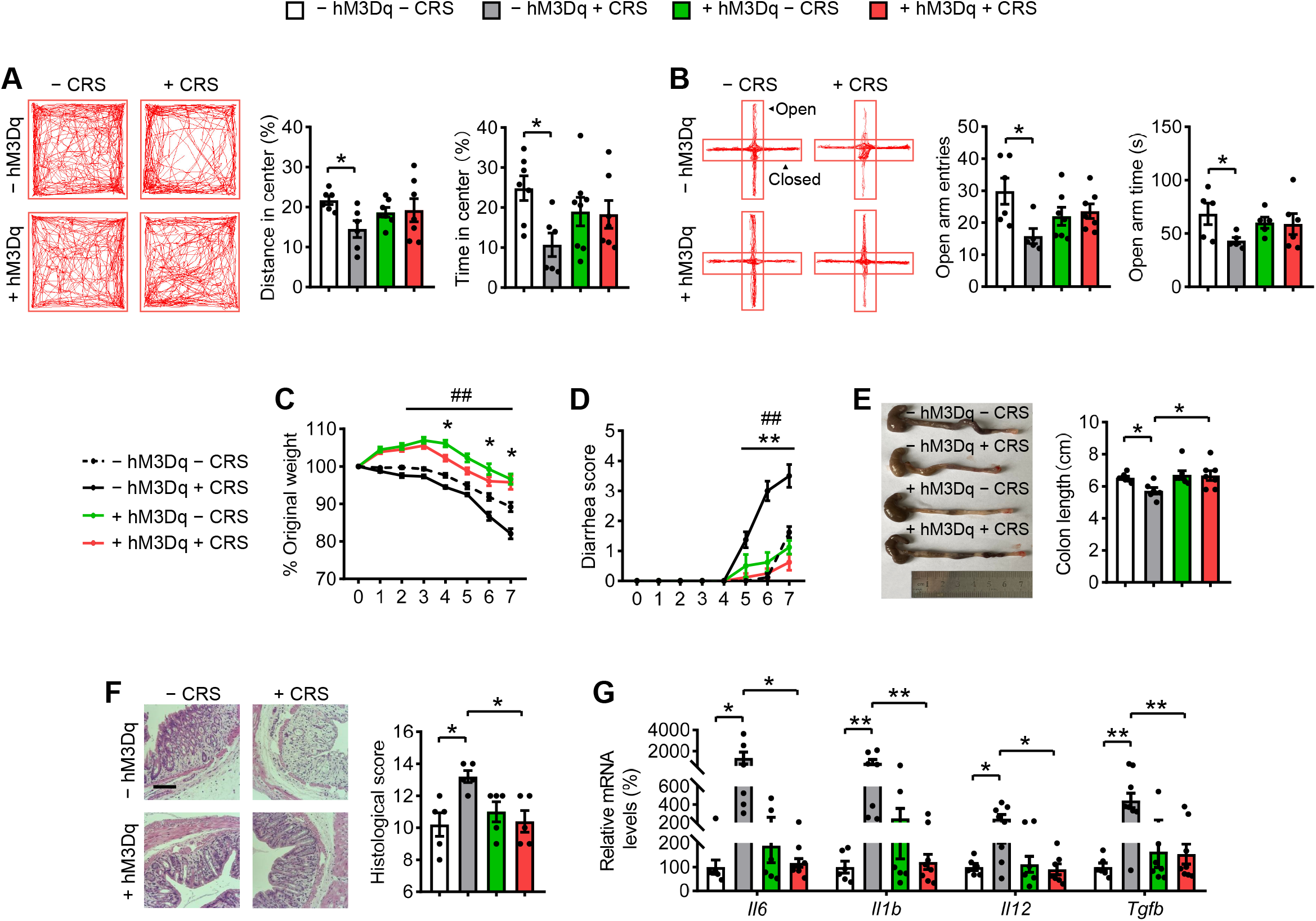
Activation of AgRP neurons reverses CRS-induced anxiety behaviors and colitis. (A) Representative tracks and statistical results in OF test. (B) Representative tracks and statistics in EPM test. (C) Percentage of body weight loss. (D) Scores of diarrhea. (E) Gross morphology and length of the colon. (F) H&E staining and histological scores of the colon tissues. Scale bar, 110 μm. (G) qRT-PCR analysis of mRNA expression of inflammatory cytokines (*Il6, Il1b, Il12*, and *Tgfb)* in the distal colon tissues. Studies for A-B were conducted using 10- to 12-week-old AgRP-Cre mice receiving AAV expressing mCherry (− hM3Dq) or hM3Dq (+ hM3Dq), all mice experienced unstressed (− CRS) or stressed (+ CRS) treatment for 14 days. Behavioral tests were performed 30 min after single CNO injection on day 15 (A) and day 16 (B). C-G were performed using - hM3Dq mice and + hM3Dq mice under treatment of 3% DSS in drinking water for 7 days to induce acute colitis with (+ CRS) or without (- CRS) stress, simultaneously receiving CNO injections every 12 hours per day. Values are expressed as means ± SEM (n = 5-8 per group), with individual data points. Data were analyzed using two-way analysis of variance, followed by Tukey’s multiple comparisons test. - hM3Dq + CRS versus - hM3Dq - CRS, *P <0.05, **P <0.01; + hM3Dq + CRS versus - hM3Dq + CRS, ^#^P <0.05, ^##^P <0.01 (C-D).

### Inhibition of AgRP Neurons Promotes Anxiety-Like Behaviors and Colitis

To further confirm the role of AgRP neurons in anxiety and colitis, we investigated the phenotypes in mice with inhibition of AgRP neurons, using an inhibitory hM4Di designer receptor exclusively activated by designer drugs (DREADDs) (Krashes et al., 2011), as reflected by the reduced c-Fos immunoreactivity in AgRP neurons (Supplementary information Fig. S4A and B). Interestingly, inhibition of AgRP neurons decreased the center distance and center time in the OF test, and the number of entries and time spent in the open arms in the EPM test, indicating increased anxiety-like behaviors (Fig. 2A and B). Moreover, mice with inhibited neuronal activity of the AgRP were more sensitive to DSS-induced colitis, characterized by a more severe weight loss, gross bleeding, shortened colon length, higher histological scores, and increased pro-inflammatory cytokine levels (*Il6, Il1b, Il12*, and *Tgfb)*, compared with control mice (Fig. 2C–G, Supplementary information Fig. S4C). Collectively, these results indicate that inhibition of AgRP neurons mimics stress-induced anxiety-like behaviors and colitis.

**Figure 2.**
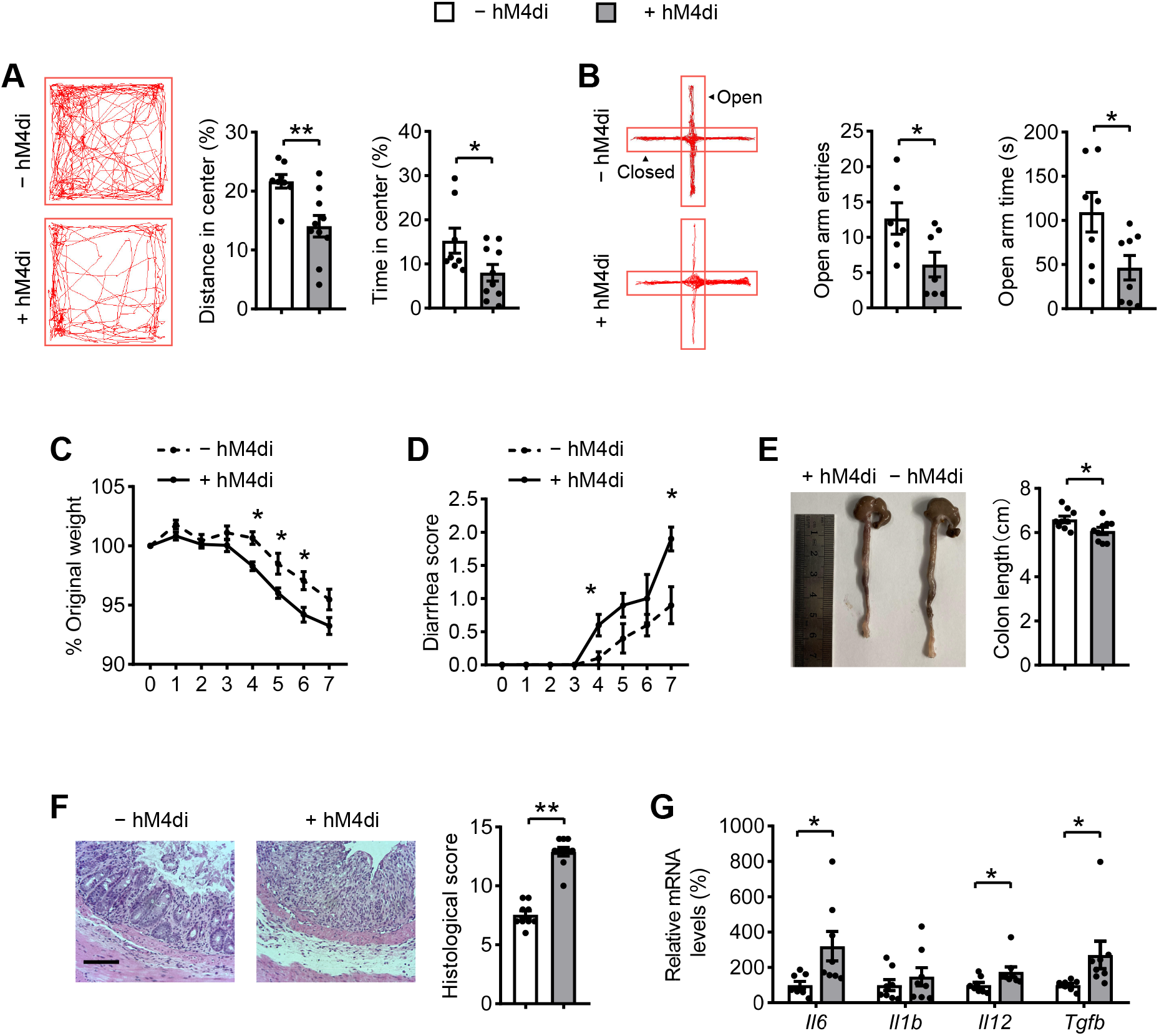
Inhibition of AgRP neurons mimics CRS-increased susceptibility to anxiety and colitis. (A) Representative tracks and statistical results in OF test. (B) Representative tracks and statistics in EPM test. (C) Percentage of body weight loss. (D) Scores of diarrhea. (E) Gross morphology and length of the colon. (F) H&E staining and histological scores of the colon tissues. Scale bar, 110 μm. (G) qRT-PCR analysis of mRNA expression of inflammatory cytokines (*Il6, Il1b, Il12*, and *Tgfb)* in the distal colon tissues. Studies for A-B were conducted using 10- to 12-week-old AgRP-Cre mice receiving AAV expressing mCherry (− hM4Di) or hM4Di (+ hM4Di), all mice received CNO injections every 12 h per day. Behavioral tests were performed 30 min after single CNO injection on day 22 (A) and day 23 (B). C-G were performed using - hM4Di mice and + hM4Di mice with 3% DSS in drinking water for 7 days to induce acute colitis after 21 days of CNO injections. Values are expressed as means ± SEM (n = 8-10 per group), with individual data points. Data were analyzed using two-tailed unpaired Student’s *t* test.

### Overexpression of c-Jun in AgRP Neurons Confers Resistance to CRS-Induced Anxiety-Like Behaviors and Colitis Susceptibility

We then explored the possible involvement of c-Jun in the CRS-induced effects and found that the activity of AgRP neurons decreased through c-Jun ablation and increased through c-Jun overexpression both *in vivo* and *in vitro* (Supplementary information Fig. S5A-D). IF staining confirmed a decrease in c-Jun protein levels in the AgRP neurons of stressed mice (Supplementary information Fig. S6A). If the reduced c-Jun expression was important under stress, the activation of c-Jun in AgRP neurons should be expected to ameliorate CRS-induced effects. To test this possibility, we overexpressed c-Jun in AgRP neurons by injecting AAVs expressing c-Jun into the AgRP-irs-Cre mice (Supplementary information Fig. S6B and C). As predicted, mice with c-Jun overexpression were resistant to CRS-induced weight loss and colon shortening (Supplementary information Fig. S6D and E). Consistently, overexpressed groups spent more time and distance in the center as evaluated in the OF test compared with the control group after CRS (Supplementary information Fig. S6F). Similarly, in the EPM test, groups with overexpressed c-Jun had more entries to the open arm (Supplementary information Fig. S6G). Stressed c-Jun-overexpressing mice were more resistant to DSS-induced body weight loss compared with stressed control mice (Supplementary information Fig. S6H and I). The diarrhea scores and colon length were also relieved in the overexpressed group (Supplementary information Fig. S6J and K). Furthermore, signs of colon colitis were markedly ameliorated in mice with overexpressed c-Jun, as evidenced by the decreased epithelial damage and lymphocyte infiltration, as well as reduced mRNA expression of inflammatory cytokines (Supplementary information Fig. S6L and M). These data indicate that overexpression of c-Jun in AgRP neurons protected the mice from CRS-induced anxiety and colitis.

### Deletion of c-Jun in AgRP Neurons Facilitates Anxiety-Like Behaviors and Colitis

To further confirm the role of c-Jun in AgRP neurons, we generated mice with c-Jun knockdown in AgRP neurons (*c-Jun*^ΔAgRP^), as confirmed by the reduced c-Jun expression in AgRP neurons (Supplementary information Fig. S7A). The corticosterone concentration was significantly higher in the sera of *c-Jun*^ΔAgRP^ mice than in control mice, suggesting increased stress (Supplementary information Fig. S7B). Moreover, the body weight of *c-Jun*^ΔAgRP^ mice was slightly lower than that of control mice (Supplementary information Fig. S7C). Although the classic function of AgRP neurons is to regulate food intake, we did not detect any difference in the change of food intake in *c-Jun*^ΔAgRP^ mice (Supplementary information Fig. S7D).

The *c-Jun*^ΔAgRP^ mice displayed obvious anxiety-like behaviors, reflected as a shorter time and less distance in the center in the OF test (Fig. 3A), and fewer entries and time in the open arms as evaluated in the EPM test (Fig. 3B). The clinical signs of colitis, including weight loss, rectal bleeding and colon shortening, were more severe in *c-Jun*^ΔAgRP^ mice than in controls after DSS treatment (Fig. 3C-E). The epithelial damage, including mucosal erosion, crypt loss, lymphocyte infiltration, and the mRNA expression of proinflammatory cytokines were also significantly increased in the colons of *c-Jun*^ΔAgRP^ mice compared with controls (Fig. 3F and G). These data indicate that deletion of *c-Jun* in AgRP neurons is sufficient to induce anxiety-like behaviors and colitis susceptibility in the absence of stress.

**Figure 3.**
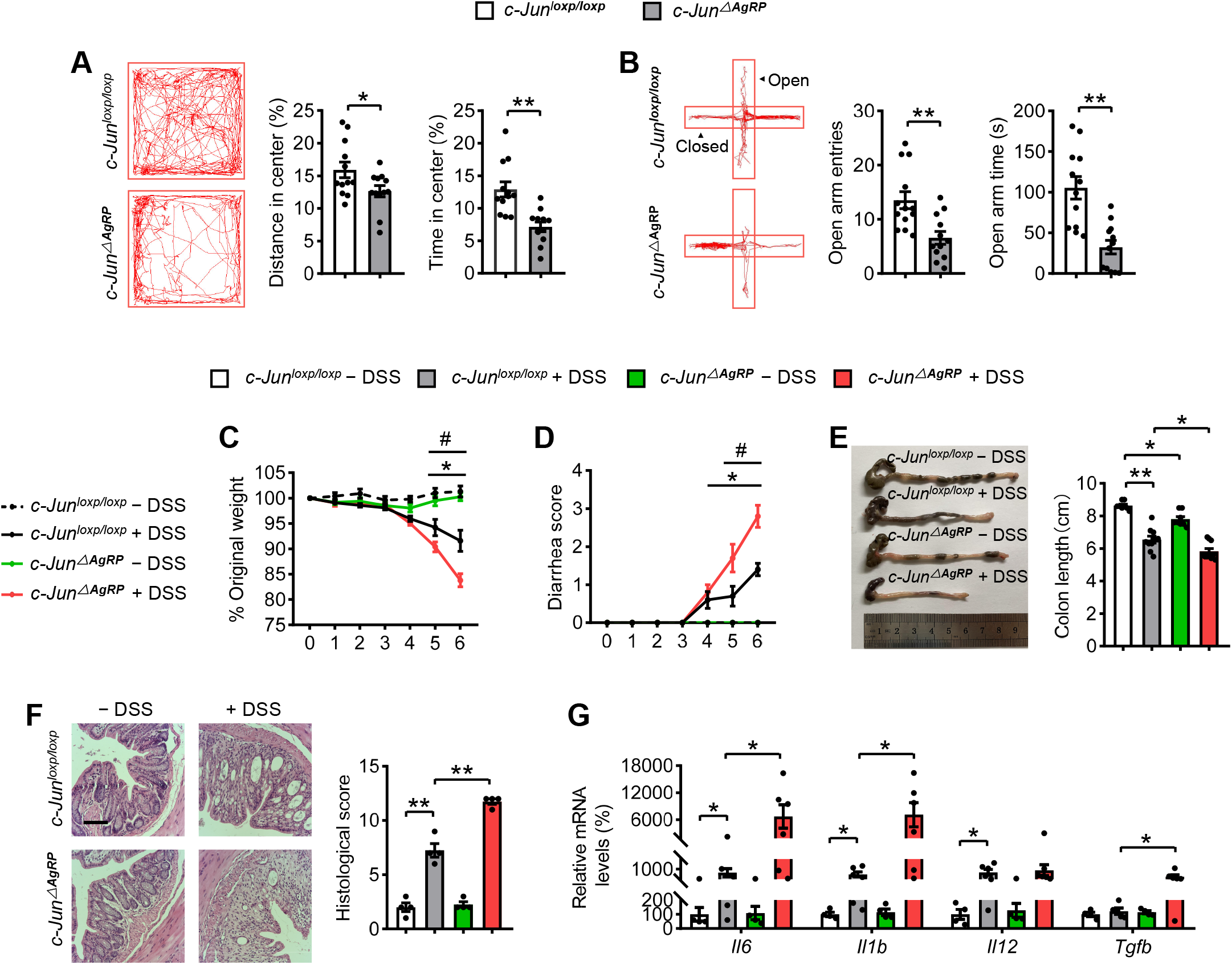
Deletion of c-Jun in AgRP neurons facilitates anxiety-like behaviors and colitis. (A) Representative tracks and statistical results in OF test. (B) Representative tracks and statistics in EPM test. (C) Percentage of body weight loss. (D) Scores of diarrhea. (E) Gross morphology and length of the colon. (F) H&E staining and histological scores of the colon tissues. Scale bar, 110 μm. (G) qRT-PCR analysis of mRNA expression of inflammatory cytokines (*Il6, Il1b, Il12*, and *Tgfb)* in the distal colonic tissues. Studies for A-B were conducted using 20- to 22-week-old control mice (*c-Jun*^loxp/loxp^) or mice with c-Jun deletion in AgRP neurons (*c-Jun*^ΔAgRP^). C-G were performed in *c-Jun*^loxp/loxp^ and *c-Jun*^ΔAgRP^ mice administrated with (+ DSS) or without (- DSS) 3% DSS for 6 days to induce acute colitis. Values are expressed as means ± SEM (n = 4-12 per group), with individual data points. Data were analyzed using two-way analysis of variance, followed by Tukey’s multiple comparisons test. *c-Jun*^loxp/loxp^ + DSS versus *c-Jun*^loxp/loxp^ - DSS, *P <0.05, **P <0.01. *c-Jun*^ΔAgRP^ + DSS versus *c-Jun*^loxp/loxp^ + DSS, ^#^P <0.05, ^##^P <0.01 (C-D).

### Relieving Anxiety with Cyamemazine (CYA) Reverses Colitis in *c-Jun*^ΔAgRP^ Mice

To investigate whether stimulated anxiety-like behaviors contributed to the increased colitis susceptibility, a typical antipsychotic drug CYA has been shown to block anxiety (Benyamina et al., 2012), was infused into the third ventricle of *c-Jun*^ΔAgRP^ mice for 7 days. The efficacy of the drug was elevated by the changed 5-HT receptor expression in the ARC (Supplementary information Fig. S8), as CYA is a potent 5-HT receptors antagonist (Benyamina et al., 2012). Interestingly, we found that treatment with CYA largely alleviated anxiety-like behaviors in *c-Jun*^ΔAgRP^ mice as demonstrated by the OF and EPM tests (Fig. 4A and B). The signs of colitis susceptibility were also alleviated in *c-Jun*^ΔAgRP^ mice, following evaluation of the examined parameters (Fig. 4C-G). These data indicate that antipsychotic drug may influence the development of colitis in mice with co-existing anxiety.

**Figure 4.**
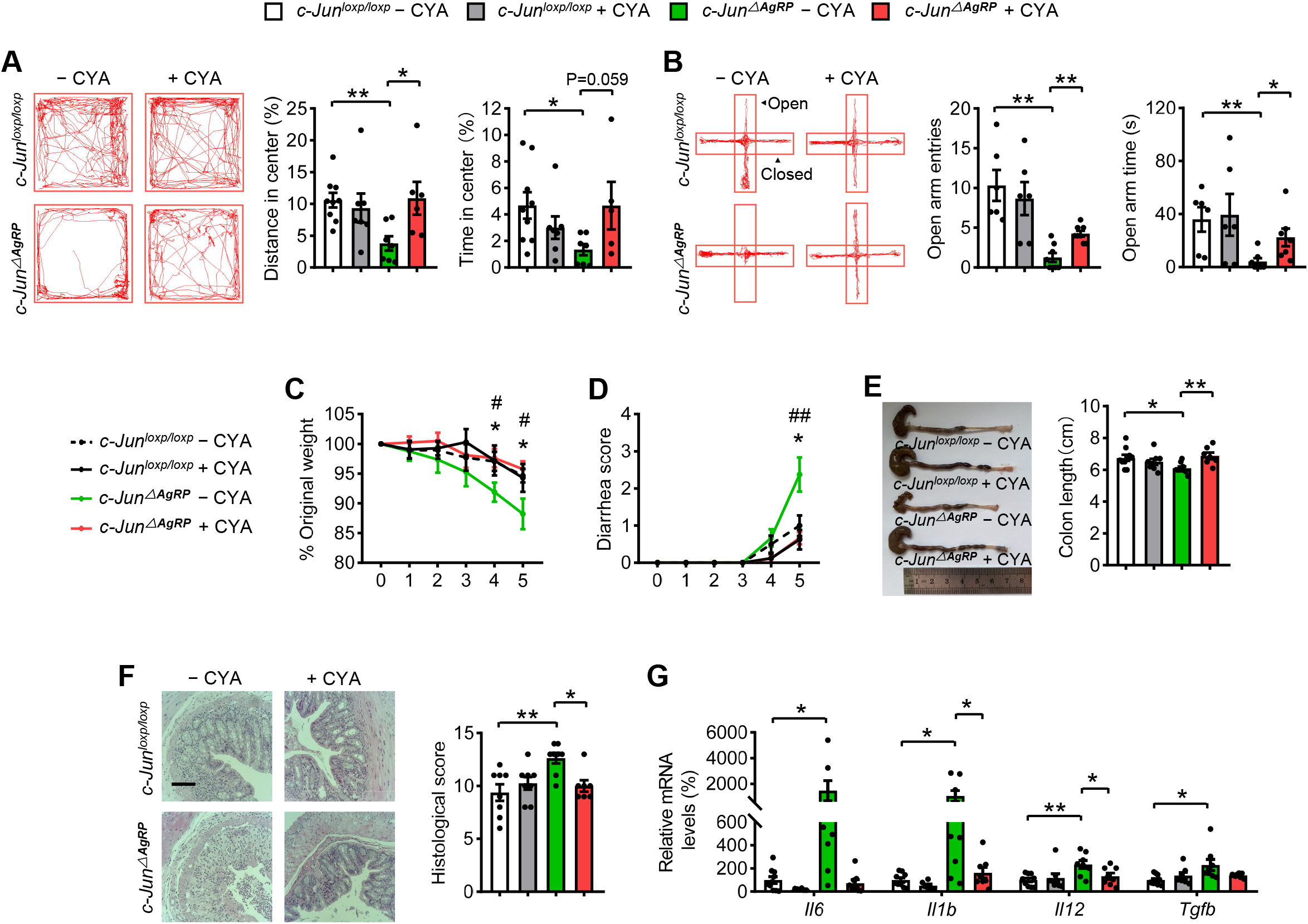
Relieving anxiety with cyamemazine (CYA) reverses colitis in *c-Jun*^ΔAgRP^ mice. (A) Representative tracks and statistical results in OF test. (B) Representative tracks and statistics in EPM test. (C) Percentage of body weight loss. (D) Scores of diarrhea. (E) Gross morphology and length of the colon. (F) H&E staining and histological scores of the colon tissues. Scale bar, 110 μm. (G) qRT-PCR analysis of mRNA expression of inflammatory cytokines (*Il6, Il1b, Il12*, and *Tgfb)* in the distal colonic tissues. Studies for A-B were conducted using 18- to 20-week-old *c-Jun*^loxp/loxp^ mice and *c-Jun*^ΔAgRP^ mice treated with (+ CYA) or without (- CYA) CYA for 7 days. Behavioral tests were performed 30 min after single CYA injection on day 8 (A) and day 9 (B). C-G were performed in *c-Jun*^loxp/loxp^ and *c-Jun*^ΔAgRP^ mice administrated with 3% DSS for 5 days to induce acute colitis with (+ CYA) or without (- CYA) CYA. Values are expressed as means ± SEM (n = 6-10 per group), with individual data points. Data were analyzed using two-way analysis of variance, followed by Tukey’s multiple comparisons test. *c-Jun*^ΔAgRP^ - CYA versus *c-Jun*^loxp/loxp^ - CYA, *P <0.05, **P <0.01. *c-Jun*^ΔAgRP^ + CYA versus *c-Jun*^ΔAgRP^-CYA, ^#^P <0.05, ^##^P <0.01 (C-D).

### The Increased Colitis Susceptibility in *c-Jun*^ΔAgRP^ mice is Mediated by THBS1

Because the brain conveys the neural, endocrine, and circulatory messages to the gut (Bonaz and Bernstein, 2013; Gracie et al., 2019; Mawdsley and Rampton, 2005), to elucidate the underlying mechanisms of the observed effects, we conducted mass spectrometry to explore the possible secreted proteins in the sera of the mice with c-Jun overexpression and control groups with or without CRS under colitis conditions. We identified 22 secreted proteins that significantly differed in abundance between the three groups (Fig. 5A and B; Supplementary information Table S2-3). Among these proteins, we focused on thrombospondin 1 (THBS1), which showed the most dramatic change and is well known for its anti-angiogenic and anti-inflammatory properties (Adams and Lawler, 2011). The secreted levels of THBS1 were significantly reduced after CRS and notably increased after c-Jun rescue in AgRP neurons (Supplementary information Fig. S9A-C). Moreover, the serum levels of THBS1 were reduced in *c-Jun*^ΔAgRP^ mice (Fig. 5C) and increased by treatment with CYA (Supplementary information Fig. S9D), indicating it may have potential role in linking anxiety and colitis. To test this hypothesis, we treated *c-Jun*^ΔAgRP^ mice with THBS1 (Bai et al., 2020) and found that it markedly alleviated colitis, as shown by the resistant effects on the corresponding changes in the body weight loss, bleeding score, colon length, colon histochemical analysis, and the expression of pro-inflammatory factors (Fig. 5D-H). These results suggest that THBS1 suppresses intestinal mucosal inflammation and may serve as a potential biomarker for stress-induced colitis.

**Figure 5.**
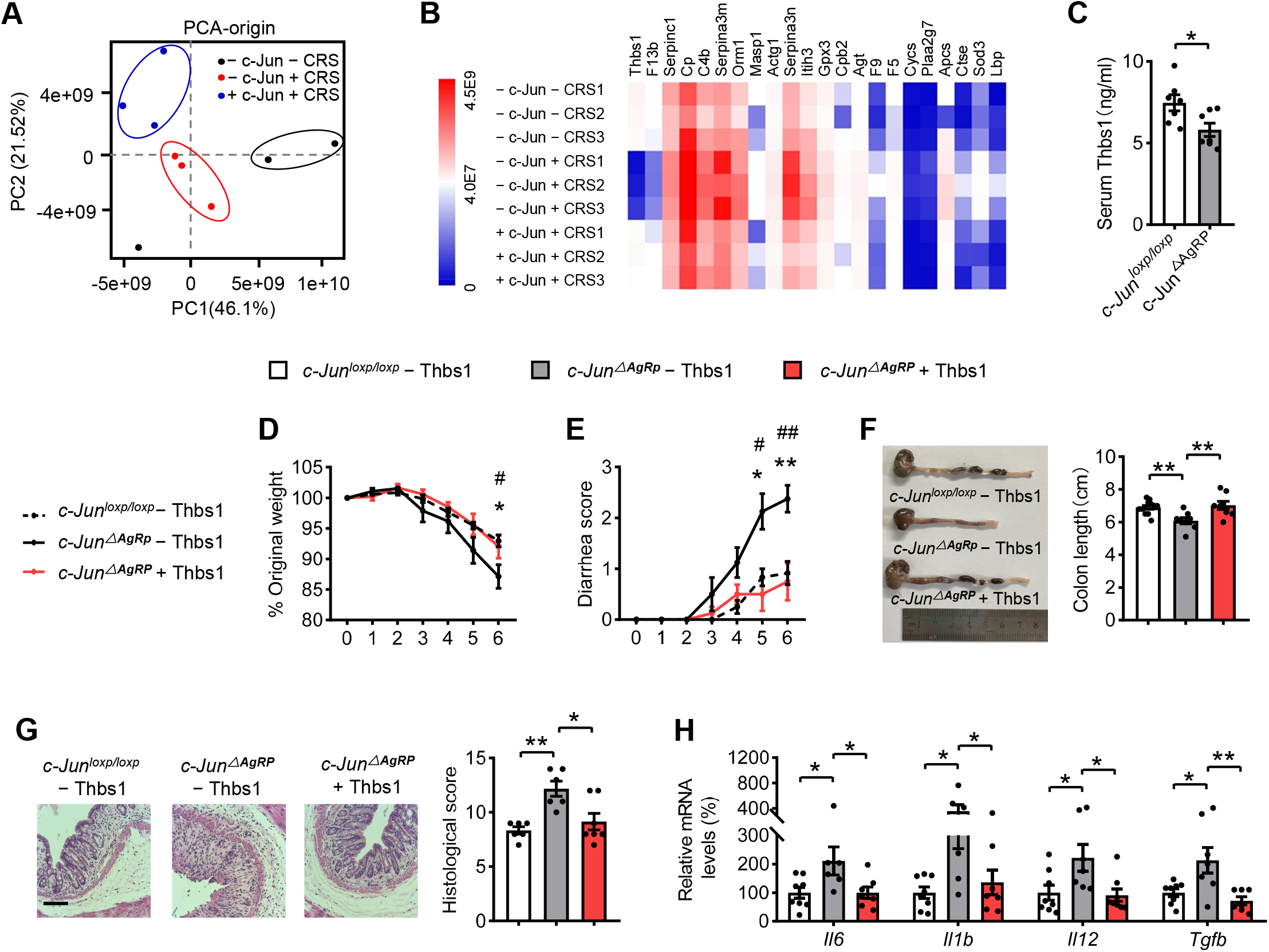
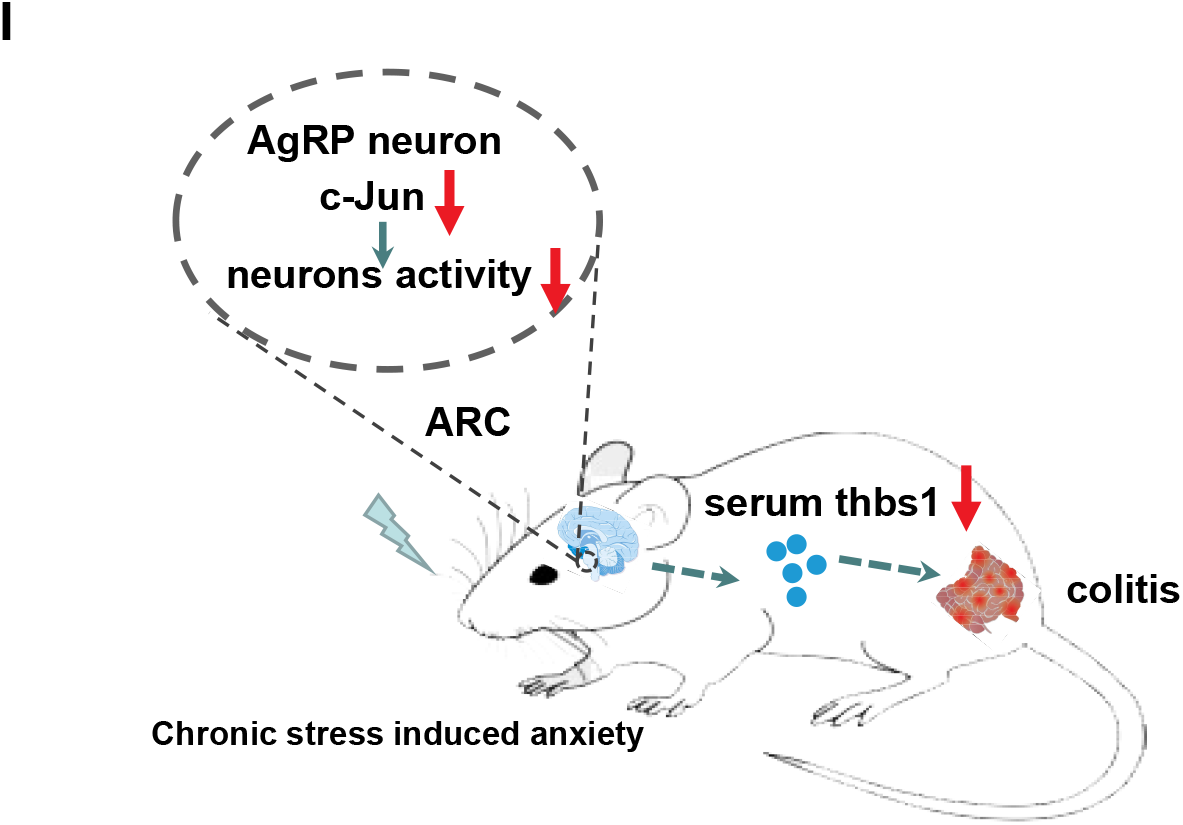
The increased colitis susceptibility in *c-Jun*^ΔAgRP^ mice is mediated by THBS1. (A) Partial least-squares discriminant analysis of protein composition. (B) Spearman’s correlation analysis of serum proteome. (C) Serum thbs1 levels. (D) Percentage of body weight loss. (E) Scores of diarrhea. (F) Gross morphology and length of the colon. (G) H&E staining and histological scores of the colon tissues. Scale bar, 110 μm. Scale bar, 110 μm. (H) qRT-PCR analysis of mRNA expression of inflammatory cytokines (*Il6, Il1b, Il12*, and *Tgfb)* in the distal colonic tissues. (I) Summary Diagram. Chronic restraint stress induces anxiety-like behaviors and colitis susceptibility, which is mediated by c-Jun in AgRP neurons. Knockdown of c-Jun in AgRP neurons decreases AgRP neurons activity and increases anxiety-like behaviors and colitis susceptibility through reducing serum THBS1 levels. Studies for A-B were conducted using - c-Jun - CRS mice, - c-Jun + CRS mice and + c-Jun + CRS mice with 3% DSS for 7 days. Serum was collected after DSS stimulation for proteomics profiling. C was conducted using 24- to 26-week-old *c-Jun*^loxp/loxp^ mice and *c-Jun*^ΔAgRP^ mice with DSS administration. D-H were conducted using 22- to 24-week-old *c-Jun*^loxp/loxp^ mice and *c-Jun*^ΔAgRP^ mice with (+ thbs1) or without (− thbs1) thbs1 supplementary, simultaneously receiving 3% DSS treatment for 6 days to induce acute colitis. Values are expressed as means ± SEM (n=3-9 per group), with individual data points. Data were analyzed using two-way analysis of variance, followed by Tukey’s multiple comparisons test. *c-Jun*^ΔAgRP^ - thbs1 versus *c-Jun*^loxp/loxp^ - thbs1, *P <0.05, **P <0.01; *c-Jun*^ΔAgRP^ + thbs1 versus *c-Jun*^ΔAgRP^ - thbs1, ^#^P <0.05, ^##^P <0.01 (D-E).

## Discussion

The brain-gut axis serves as a circuit that incorporates the state of mind and gut signals that ultimately determine the intestinal function (Bonaz and Bernstein, 2013; Gracie et al., 2019; Mawdsley and Rampton, 2005; Wu et al., 2014). Therefore, the changes of brain functions are closely related to gut metabolism abnormalities (Bonaz and Bernstein, 2013; Lee et al., 2021; Mawdsley and Rampton, 2005). Accumulating evidence indicates that mood disorders, such as anxiety or depression, often co-occur with IBD (Blackwell et al., 2021; Gracie et al., 2018; Koloski et al., 2012; Kurina et al., 2001). Several brain areas, including the hypothalamus, hippocampus and amygdala, are involved in anxiety-related behaviors (Adhikari et al., 2010; Bains et al., 2015; Liu et al., 2020). The AgRP neurons in the hypothalamic ARC particularly have gained much attention since more important functions of this neuron were discovered, such as feeding, pain sensation and depression-related behaviors (Alhadeff et al., 2018; Bell et al., 2005; Fang et al., 2021). However, a role of AgRP neurons in the comorbidity of anxiety and colitis has not been reported.

In current study, we used CRS model and chemogenic strategy to investigate the possible involvement of AgRP neurons in stress-induced anxiety and colitis. We found that AgRP neuronal activity was inhibited by CRS. However, in another study under stressed model, AgRP neurons were not affected (Qu et al., 2020). The different response may be attributable to the variation in the duration of treating the mice (4 hours per day in our work versus 1 hour per day in their study), which is likely to result in different levels of stress. The significance of the inhibited AgRP neurons in CRS was further confirmed by gain- and loss-of AgRP neuron function experiments. To our knowledge, this is the first to demonstrate that convergent regulation of colitis and stress is integrated in the AgRP neurons in the ARC, providing important evidence that psychiatric disorders, such as anxiety, may influence colitis through central neuronal activity in the ARC. Moreover, though some mechanisms are proposed for IBD (Molodecky et al., 2012), the neuronal signals are largely unknown. Our results provide new perspective for understanding the neuronal regulation for IBD. In addition, another study shows that inhibition of AgRP neuron causes depression (Fang et al., 2021), another type of psychiatric disorder (de Kloet et al., 2005), suggesting that AgRP neurons might also play an important role in other psychiatric disorders, as well as diseases related to them. As a number of studies have shown that AgRP neurons facilitate food-seeking behavior (Aponte et al., 2011; Krashes et al., 2011), it remains to be tested whether the role of Agrp neurons in anxiety-like behaviors is influenced by the behaviors driven by feeding. However, the increased food intake did not contribute to the improvement of colitis as evaluated by pair-fed experiments in current study.

We then explored the specific neural molecules regulating stress-induced anxiety and colitis. We found that the expression of the stress-related gene c-Jun was decreased in AgRP neurons under CRS conditions and that knockdown of c-Jun in AgRP neurons mimicked the effect of CRS but reversed these after overexpression of c-Jun. These results demonstrate a novel regulatory role of the central c-Jun in stress-induced anxiety and colitis and provide a potential therapeutic approach for the treatment of these diseases. However, it is unclear how c-Jun inhibited neuronal activity under stress. Considering that c-Jun plays a role in the regulation of neuron injury and in the promotion of axon regeneration (Raivich et al., 2004), we speculated that stress may lead to the damage of AgRP neurons, thus leading to a change in neuronal activity. However, this possibility requires further investigation.

Moreover, we provided important evidence that anxiety may influence colitis susceptibility by using anxiolytic drug, as well as the possibility for the treatment of IBD with co-existing anxiety. To gain further insights into how anxiety influences colitis, we performed proteomic analysis and serum measurement and found that the levels of secreted protein THBS1 were decreased by CRS in a c-Jun dependent manner, suggesting that it may function as downstream in linking c-Jun regulated anxiety and colitis. Thrombospondins are a family of extracellular matrix proteins, which were first identified in platelets stimulated with thrombin (Baenziger et al., 1971). After treatment with DSS, THBS1-deficient mice show a higher level of crypt damage and deeper lesions, which are reversed by treatment with a THBS1 mimetic peptide (Punekar et al., 2008). The importance of THBS1 in mediating anxiety-associated colitis was confirmed by the reversal effect of recombinant THBS1 on colitis in *c-Jun*^ΔAgRP^ mice. Because THBS1 levels were correspondingly changed with the status of anxiety and colitis, suggesting that it might be used as a biomarker for the comorbidity of these diseases. However, it remains unclear for the source of secreted THSB1 protein in the sera, as THBS1 can be produced and secreted into the extracellular space of many cell types, including the activated endothelium, intestinal epithelial cells, and astrocytes (Adams and Lawler, 2011; Christopherson et al., 2005; Fang et al., 2015). In addition, the sympathetic nerve and the vagus nerve, which have been shown to be important mediators of central nervous system outputs to the peripheral tissues (Bonaz and Bernstein, 2013; Ghia et al., 2008), may also be involved in the regulation of anxiety-promoted colitis susceptibility. These questions remain for future investigation.

Interestingly, we found that chemogenic manipulation or c-Jun knockdown-induced inhibition of AgRP neurons promotes anxiety-like behaviors and colitis in the absence of stressor, suggesting that signals inhibited AgRP neurons may be potential target for the treatment of anxiety and/or colitis. Therefore, our results are also important for understanding the mechanisms for those without obvious stress but present with the comorbidity of psychiatric disorders and IBD.

In summary, our present findings revealed that AgRP neuronal activity in the ARC is an important link between anxiety-like behavior and intestinal inflammation (Fig. 5I). The importance of these findings is that we have uncovered the specific neurons and signals in the brain underlying the regulation of the anxiety and colitis comorbidity. Our results provide evidence that CRS-induced anxiety and colitis is mediated through an unexpected neurons AgRP neurons. Moreover, we demonstrated c-Jun as a target in AgRP neuron for stress-induced anxiety and colitis. Furthermore, we identified the secreted protein Thbs1 function in linking anxiety and colitis and as a biomarker for anxiety-colitis comorbidity. These results provide a new perspective for exploring the brain in the regulation of intestinal inflammation homeostasis, and further provide a new central target for the therapeutic intervention of stress-induced psychiatric disorders and intestinal metabolism dysfunction.

## Materials and Methods

### Mice and Treatment

Adult male C57BL/6 wild-type (WT) mice were purchased from Shanghai Laboratory Animal Co., Ltd. (Shanghai, China). *c-Jun*^loxp/loxp^ mice (generously provided by Dr. Erwin F. Wagner, Cancer Cell Biology Program, Spanish National Cancer Research Center) were crossed with mice expressing Cre recombinase under control of the AgRP promoter to generate *c-Jun*^ΔAgRP^ mice. The efficiency of AgRP-specific *c-Jun* deletion was evaluated by mating Ai9 (tdTomato) reporter mice (Madisen et al., 2010) with transgenic mice expressing Cre under control of the AgRP promoter (AgRP-irs-Cre mice), both obtained from Jackson Laboratory (Bar Harbor, ME, USA). Transgenic mice and their littermates were used in experiments at the indicated ages. Mice were subcutaneously injected with recombinant human thrombospondin 1 (THBS1) protein (0.5 mg/kg per day; Novoprotein Scientific Inc., Shanghai, China) (Bai et al., 2020) or vehicle (phosphate-buffered saline) for the indicated period. To study the effect of anxiolytic drug on anxiety and colitis, cyamemazine (0.25 ug/side, MCE) (Xia et al., 2021) were i.c.v. administrated.

Mice were maintained under controlled temperature (23°C), humidity (50–60%), and illumination (12-h light/12-h dark cycle), and provided *ad libitum* access to food and water. All animal experiments were conducted in accordance with the guidelines of the Institutional Animal Care and Use Committee of Shanghai Institute for Nutritional Sciences, Chinese Academy of Sciences.

### Chronic Restraint Stress (CRS) Model

The CRS mouse model was performed as described previously (Gao et al., 2018; Liu et al., 2020). In brief, the mice were individually placed in a 50-mL polypropylene conical tube with multiple holes for ventilation and were restrained to prevent back-and-forth movement. Restraint was applied for 4 h per day from 10:00 a.m. to 2:00 p.m. for the number of days indicated.

### Colitis Model Establishment

To establish the dextran sulfate sodium (DSS)-induced colitis model, the drinking water of the mice was supplemented with 3% (w/v) DSS (40 kDa; Aladdin, Shanghai, China) as described previously (Tian et al., 2019), and the colon length was determined at the end of the experiments. Diarrhea scores were assessed as described previously (Rachmilewitz et al., 2002).

### Stereotaxic Surgery and Viral Injections

Surgery was performed as reported previously (Yuan et al., 2020) with a stereotaxic frame (Stoelting, Wood Dale, IL, USA). Viral injection coordinates (in mm, midline, bregma, dorsal surface) are as follows: for ARC (± 0.3, 1.5, 5.9), for the third ventricle (0, 1.5, 5.6) (Deng et al., 2017; Xia et al., 2021). To rescue the expression of *c-Jun* specifically localized in AgRP neurons, AgRP-irs-Cre mice were bilaterally injected with 300 nL of a Cre-dependent adeno-associated virus (AAV) vector containing *c-Jun* in the opposite orientation flanked by two inverted loxP sites [AAV9-EF1a-DIO-c-Jun-mCherry, 2.5 x 10^12^ particle-forming units (PFU)/mL] into the ARC, or with an AAV vector containing only mCherry in the opposite orientation flanked by two inverted loxP sites (AAV9-EF1a-DIO-mCherry, 2.5 x 10^12^ PFU/mL) as controls.

### Designer Receptor Exclusively Activated by Designer Drugs (DREADDs)

To inhibit AgRP neuronal activity, AgRP-irs-Cre mice were stereotaxically injected with 300 nL of a Cre-dependent AAV encoding an inhibitory DREADD GPCR (hM4Di) (AAV9-EF1a-DIO-hM4Di-mCherry, 8 x 10^12^ PFU/mL) or an AAV encoding only mCherry (AAV9-EF1a-DIO-mCherry, 7 x 10^12^ Pfu/mL) as controls, bilaterally into the ARC.

For chemogenetic activation of AgRP neurons, AgRP-irs-Cre mice were stereotaxically injected with 300 nL of a Cre-dependent AAV encoding an excitatory DREADD GPCR (hM3Dq) (AAV9-EF1a-DIO-hM3D(Gq)-mCherry, 3 x 10^12^ PFU/mL) or an AAV encoding only mCherry (AAV9-EF1a-DIO-mCherry, 3 x 10^12^ PFU/mL) as controls, bilaterally into the ARC. After 3 weeks of recovery, all mice were then intraperitoneally injected with clozapine N-oxide (CNO) (MedChemExpress, NJ, USA) at 0.3 mg/kg of body weight for hM4Di (Krashes et al., 2011) silencing and for hM3Dq activation (Krashes *et al*., 2013) every 12 h for indicated days.

### Isolation and Treatment of Primary Hypothalamic Neurons

The mouse primary cultures of hypothalamic neurons were referred as previously described (Deng et al., 2018). sh-c-Jun and c-Jun over-expressed (pCMV-c-Jun) plasmid were transfected into cells using lipofectamine 3000 reagent (Invitrogen; Carlsbad, CA, USA) according to the manufacturer’s recommendation. The shRNA sequence for mouse c-Jun was 5’-GCTAACGCAGCAGTTGCAAAC-3’.

### Open Field (OF) Test

The OF test was performed as in previous studies (Fan et al., 2019; Liu et al., 2020). In brief, a white open field box (50 x 50 x 50 cm; length x width x height) was divided into a center field (25 x 25 cm) and a periphery field for analysis purposes. The track was analyzed using LabState (AniLab) by recognizing the central body point of the mouse throughout a 10-min session. Less time and locomotion spent in the center of the box were interpreted as anxiety-like behaviors.

### Elevated Plus Maze (EPM) Test

The EPM test was performed as previously described (Fan et al., 2019; Liu et al., 2020). The elevated plus maze made of plastic and consisted of two white open arms without walls and two white enclosed arms with walls (25-cm long, 5-cm wide, 15-cm high). The maze was placed 60 cm above the floor. Mice were introduced into the center quadrant with their back facing an open arm. The ANY-maze video tracking system (Anilab) was used to track and analyze the time of mice spent in the open arms and their entries into the open arms throughout a 10-min session. Anxiety was evaluated by fewer movements into the open arms and less time spent there.

### RNA Isolation and Quantitative Real-time (qRT)-PCR

RNA extraction and qRT-PCR were performed as described previously (Deng et al., 2018). The primer sequences used in this study are provided in Supplementary information, Table S1

### Histological Scoring

Histological scoring was performed as described previously (Hu et al., 2019). Hematoxylin and eosin (H&E)-staining of colonic tissue sections were scored in a blinded fashion for determining the degree of inflammation and tissue damage on separate scales from 0 to 6.

### Serum Measurements

The proteomics was performed by the serum after removing common high-abundance protein through using a Thermo Fisher’s serum High Abundance protein removal reag ent (High Select™ Top14 Abundant Protein Depletion Mini Spin Columns, A36370, Thermo fisher, US). The mass spectrometer Thermo Scientific Q Exactive was perfor med for Label-free quantification detection and data was analyzed by Proteome Discoverer 2.2. The secreted proteins with fold change (FC) ⩾1.5 and p-value < 0.05 were considered to be differential proteins. Volcano plots were used to filter the proteins of interest which based on log_2_(FC) and −log_10_(P-value) of the secreted proteins (Zhu et al., 2011). The raw data is available in the link: https://datadryad.org/stash/share/20xmGKT2z--QNG0ebHfHU8FR8Vy96LuKXe4AhPwHWc4. Serum corticosterone levels were measured with an enzyme-linked immunosorbent assay (ELISA) kit (ADI-901-0 97; Enzo Life Science, Farmingdale, NY, USA) as described previously (Lee et al., 20 20). The THBS1 levels were measured using an ELISA kit (mlbio, Shanghai, China) according to the manufacturer’s recommendations.

### Immunofluorescence (IF) Staining

Immunofluorescence staining was performed as described previously (Yuan et al., 2020) with primary antibodies to c-Jun (1:1000, Cell Signaling Technology, Danvers, MA, USA) and c-Fos (1:1000, Cell Signaling Technology, mAb2250 or 1:500, Santa Cruz Biotechnology, Santa Cruz, CA, USA). c-Fos staining was coupled with a TSA Plus Fluorescein KIT (NEL741001KT, Perkinelmer, Waltham, MA, USA).

### Statistical Analysis

Experimental data are expressed as the mean ± standard error of the mean (SEM) of the number of tests stated for each experiment. Statistical comparisons were made using either two-tailed Student’s *t* test or two-way analysis of variance, followed by Tukey’s multiple comparisons test, as indicated in the figure legends. All statistical tests were performed using GraphPad Prism, version 8.0 (GraphPad Software, San Diego, CA, USA). In addition, the individual data points on each graph were shown in order to reflect the individual variability of the measures. *P <0.05, **P <0.01.

## Acknowledgements

We would like to thank Dr. Erwin F. Wagner (Cancer Cell Biology Program, Spanish National Cancer Research Center) and Lijian Hui (Chinese Academy of Sciences Center for Excellence in Molecular Cell Science) for providing c-Jun loxp/loxp mice.

## Author Contributions

Fuxin Jiao, Zhanju Liu and Feifan Guo designed the project and analyzed the data; Fuxin Jiao, Xiaoming Hu, Hanrui Yin, Feixiang Yuan, Ziheng Zhou performed the experiments; Fuxin Jiao, Zhanju Liu, and Feifan Guo wrote the manuscript. Wei Wu and Shanghai Chen provided experimental materials; All authors discussed and revised the manuscript.

## Supplementary Figures and Figure Legends

**Supplementary Figure 1.**
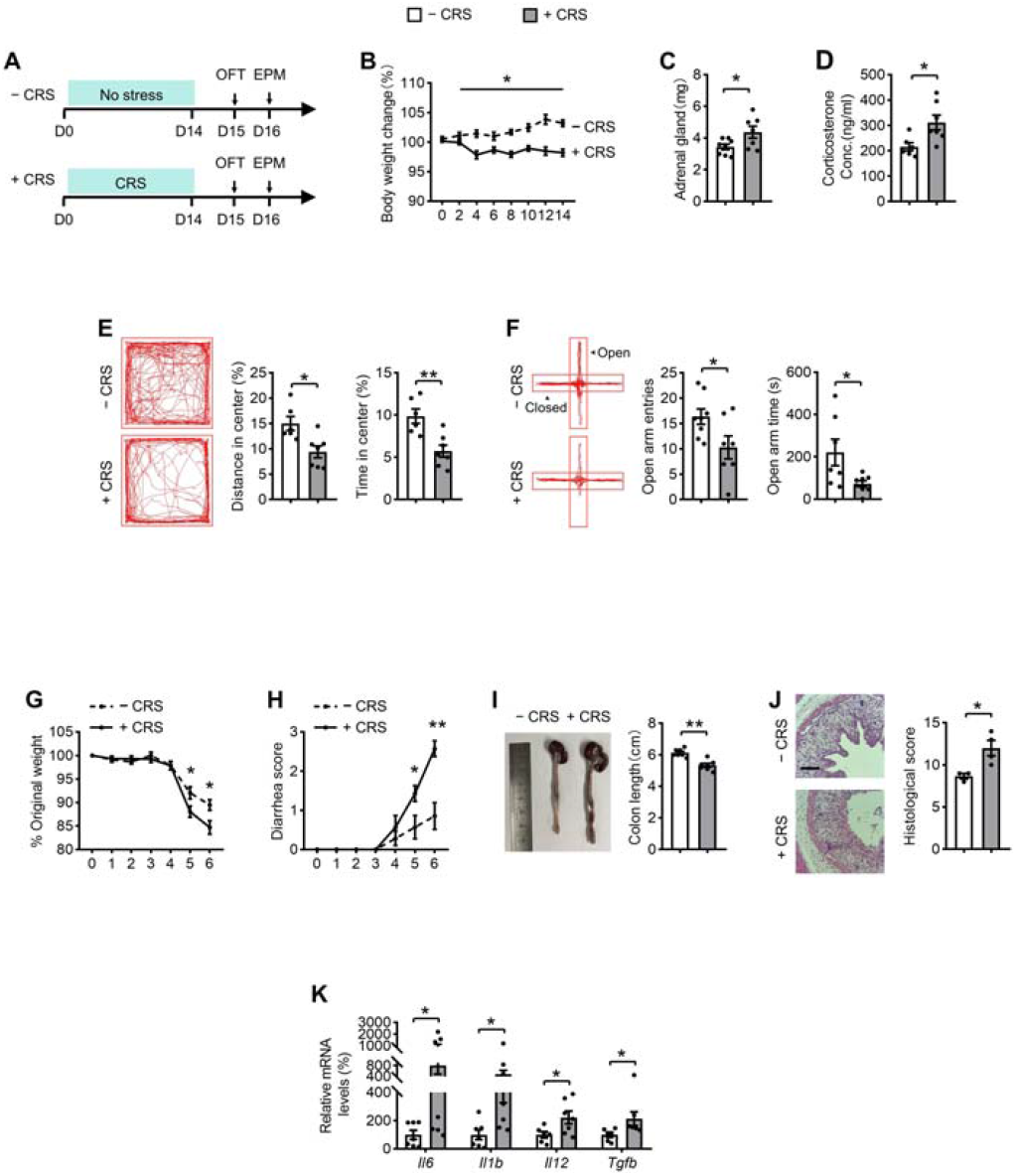
Chronic restraint stress (CRS) induces anxiety-like behaviors and increases the susceptibility to colitis. (A) Schematic showing the CRS experimental protocol. (B) Changes of body weight. (C) Adrenal gland weights. (D) Serum corticosterone levels. (E) Representative tracks and statistical results in OF test. (F) Representative tracks and statistics in EPM test. (G) Percentage of body weight loss. (H) Scores of diarrhea. (I) Gross morphology and length of the colon. (J) H&E staining and histological scores of the colon tissues. Scale bar, 110 μm. (K) qRT-PCR analysis of mRNA expression of inflammatory cytokines (*Il6, Il1b, Il12*, and *Tgfb)* in the distal colon tissues. Studies for A-F were conducted using 12-week-old WT mice with unstress (- CRS) or stressed (+ CRS) treatment for 14 days. Behavioral tests were performed on day 15 (E) and day 16 (F). G-K were conducted using - CRS mice and + CRS mice with 3% DSS in drinking water for 6 days to induce acute colitis; Values are expressed as means ± SEM (n = 3–8 per group), with individual data points. Data were analyzed using two-tailed unpaired Student’s *t* test.

**Supplementary Figure 2.**
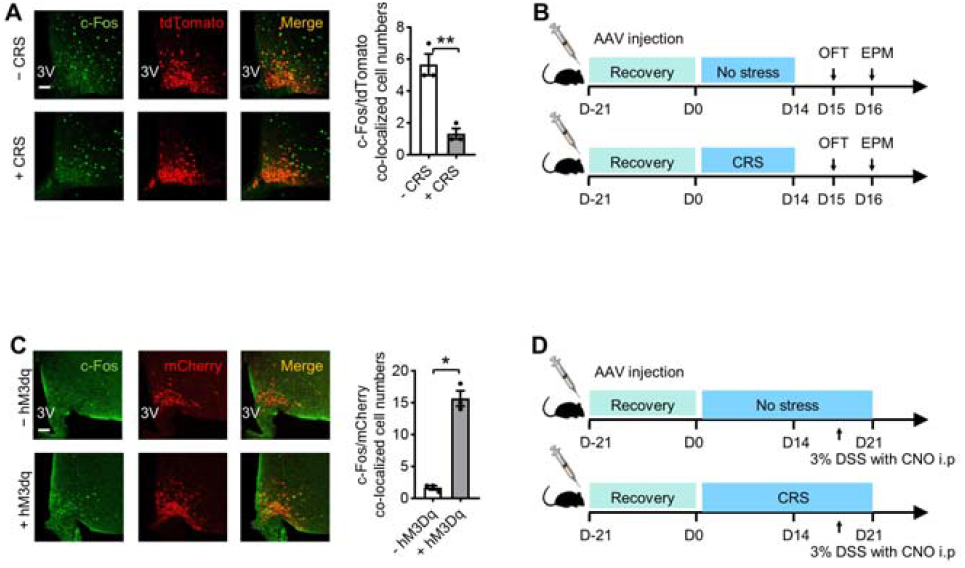
Parameters related to mice with activation of AgRP neuronal activity. (A) Immunofluorescence (IF) staining for tdTomato (red), c-Fos (green), and merge (yellow) in the ARC sections (left), and quantification of c-Fos and tdTomato co-localized cell numbers (right). Scale bar, 50 μm. c-Fos staining was coupled with a TSA Plus Fluorescein KIT. (B) Experimental timeline for CRS. (C) IF staining for mCherry (red), c-Fos (green) and merge (yellow) in ARC sections (left), and quantification of c-Fos and mCherry colocalized cell numbers (right). (D) Experimental timeline for DSS administration. Study for A was conducted using 14-week-old AgRP-Cre-Ai9 mice with (+ CRS) or without (- CRS) 14 days of stress. C was conducted using 10- to 12-week-old AgRP-Cre mice receiving AAV expressing mCherry (- hM3Dq) or hM3Dq (+ hM3Dq), both treated with one injection of CNO and 30 min later for immunofluorescence analysis. Values are expressed as means ± SEM (n = 3-4 per group), with individual data points. Data were analyzed using two-tailed unpaired Student’s *t* test.

**Supplementary Figure 3.**
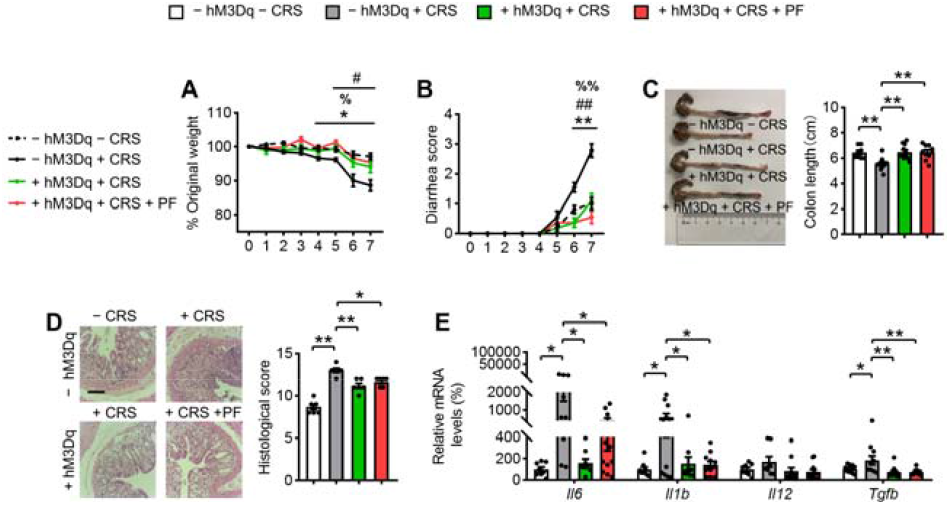
Pair-feeding has no beneficial effect on the improvement of colitis. (A) Percentage of body weight loss. (B) Scores of diarrhea. (C) Gross morphology and length of the colon. (D) H&E staining and histological scores of the colon tissues. Scale bar, 110 μm. (E) qRT-PCR analysis of mRNA expression of inflammatory cytokines (*Il16, Il1b, Il12*, and *Tgfb)* in the distal colonic tissues. Studies were conducted using 10- to 12-week-old AgRP-Cre mice receiving AAV expressing mCherry (- hM3Dq) or hM3Dq (+ hM3Dq), all mice experienced unstressed (- CRS) or stressed (+ CRS) treatment for 14 days, after that was under treatment of 3% DSS in drinking water for 7 days to induce acute colitis, simultaneously receiving CNO injections every 12 hours per day. Pair-fed (PF) experiment was administered during CNO injection. For pair-fed groups (+ hM3Dq + CRS + PF), mice were given the same diet as the - hM3Dq + CRS groups. Values are expressed as means ± SEM (n=6-11 per group), with individual data points. Data were analyzed using two-way ANOVA, followed by Tukey’s multiple comparisons test. - hM3Dq + CRS versus - hM3Dq – CRS, *P <0.05, **P <0.01; + hM3Dq + CRS versus - hM3Dq + CRS, #P <0.05, ##P <0.01; + hM3Dq + CRS + PF versus - hM3Dq + CRS, %P <0.05, %%P <0.01 (A-B).

**Supplementary Figure 4.**
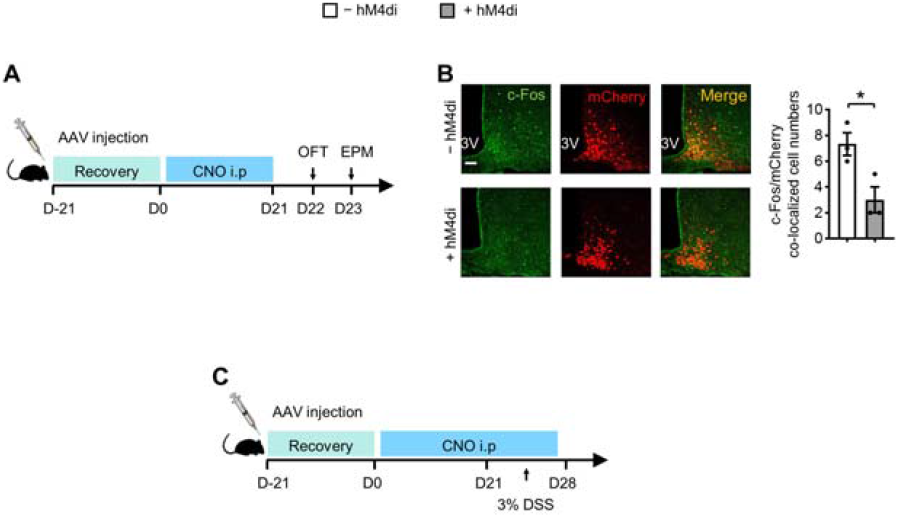
Parameters related to mice with inhibition of AgRP neuronal activity. (A) Experimental timeline for CRS. (B) Immunofluorescence (IF) staining for mCherry (red), c-Fos (green) and merge (yellow) in ARC sections (left), and quantification of c-Fos and mCherry colocalized cell numbers (right). (C) Experimental timeline for DSS administration. Study for B was conducted using 10- to 12-week-old AgRP-Cre mice receiving AAV expressing mCherry (− hM4Di) or hM4Di (+ hM4Di) both treated with one injection of CNO and 30 min later for immunofluorescence analysis. Values are expressed as means ± SEM (n = 3 per group), with individual data points. Data were analyzed using two-tailed unpaired Student’s *t* test.

**Supplementary Figure 5.**
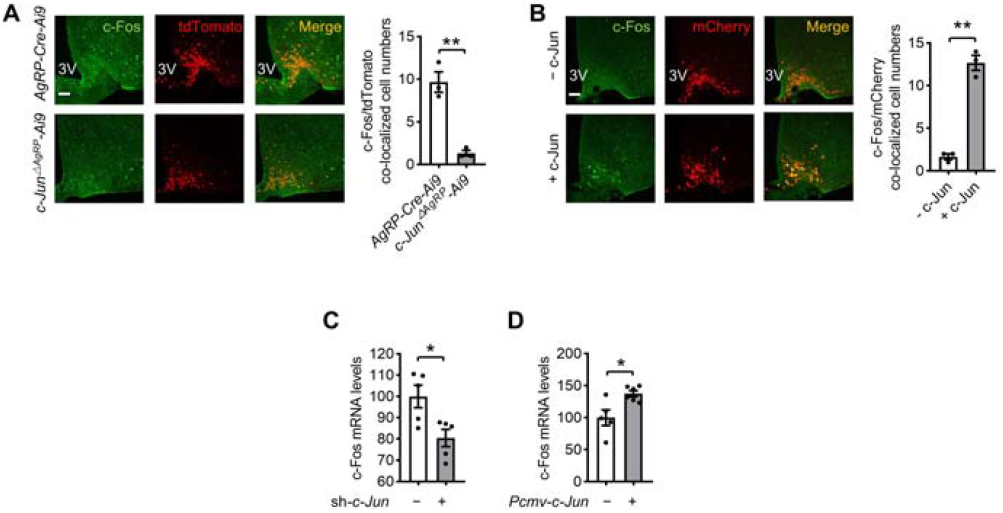
The changes of c-Fos after c-Jun deletion and overexpression in vitro and in vivo. (A) IF staining for tdTomato (red), c-Fos (green) and merge (yellow) in ARC sections (left), and quantification of c-Fos and tdTomato colocalized cell numbers (right). (B) IF staining for mCherry (red), c-Fos (green) and merge (yellow) in ARC sections (left), and quantification of c-Fos and mCherry colocalized cell numbers (right). (C-D) C-Fos mRNA levels. Study for A was conducted by mating Ai9 (tdTomato) mice with AgRP-irs-Cre mice to obtain AgRP-Ai9 mice as controls, mating Ai9 (tdTomato) mice with mice with c-Jun deletion in AgRP neurons (*c-Jun*^ΔAgRP^) to obtain *c-Jun^ΔAgRP^*-Ai9 mice. B was conducted using 12- to 14-week-old AgRP-Cre mice receiving AAV expressing mCherry (- c-Jun) or c-Jun (+ c-Jun). C was conducted on primary hypothalamus isolated from newborn mice, receiving control (*-sh-cJun*) or *sh-cJun* (+ sh-cJun) transfected with Lipo 3000. D was conducted on primary hypothalamus isolated from newborn mice, receiving control (- *pcmv-c-Jun*) or c-Jun (+ *pcmv-c-Jun)* transfected with Lipo 3000. Values are expressed as means ± SEM (n=3-6 per group), with individual data points. Data were analyzed using two-tailed unpaired Student’s *t* test.

**Supplementary Figure 6.**
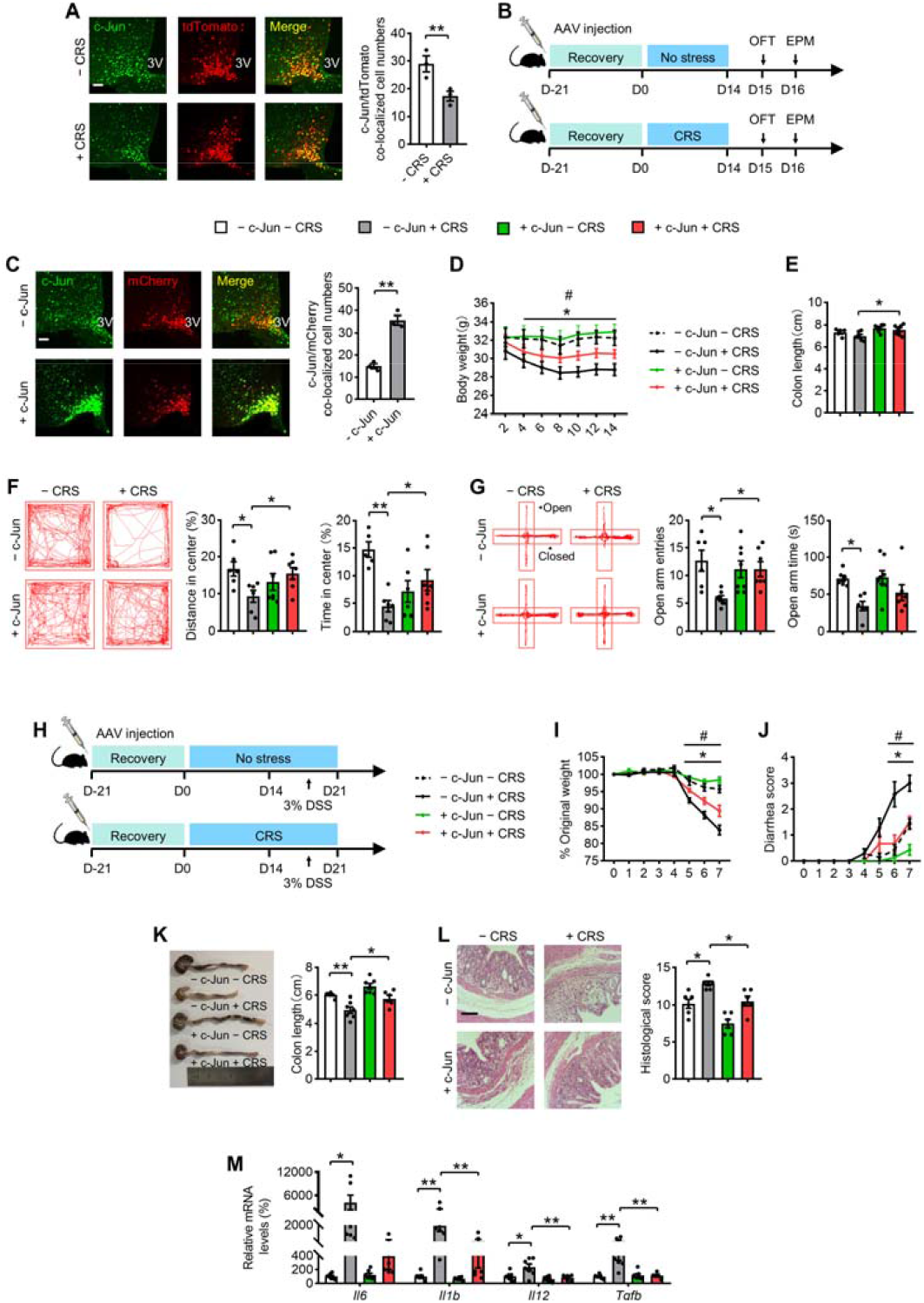
Overexpression of c-Jun in AgRP neurons leads to resistance to CRS-induced anxiety-like behaviors and colitis. (A) IF staining for tdTomato (red), c-Jun (green) and merge (yellow) in ARC sections (left), and quantification of c-Jun and tdTomato colocalized cell numbers (right). (B) Experimental timeline for CRS. (C) IF staining for mCherry (red), c-Jun (green) and merge (yellow) in ARC sections (left), and quantification of c-Jun and mCherry colocalized cell numbers (right). (D) Body weight. (E) Colon length. (F) Representative tracks and statistical results in OF test. (G) Representative tracks and statistics in EPM test. (H) Experimental timeline for DSS administration. (I) Percentage of body weight loss. (J) Scores of diarrhea. (K) Gross morphology and length of the colon. (L) H&E staining and histological scores of the colon tissues. Scale bar, 110 μm. (M) qRT-PCR analysis of mRNA expression of inflammatory cytokines (*Il16, Il1b, Il12*, and *Tgfb)* in the distal colonic tissues. Study for A was conducted using 14-week-old AgRP-Cre-Ai9 mice with (+ CRS) or without (- CRS) 14 days of stress. C was conducted using 12- to 14-week-old AgRP-Cre mice receiving AAV expressing mCherry (− c-Jun) or c-Jun (+ c-Jun), IF staining performed after 3 weeks from AAV recovery. D-G were conducted using – c-Jun mice and + c-Jun mice, both experienced unstressed (− CRS) or stressed (+ CRS) treatment for 14 days. Behavioral tests were performed on day 15 (F) and day 16 (G). H-M were conducted using - c-Jun mice and + c-Jun mice receiving 3% DSS in drinking water for 7 days to induce acute colitis, after stress (+ CRS) or unstress (− CRS). Values are expressed as means ± SEM (n=3-8 per group), with individual data points. Data were analyzed using two-tailed unpaired Student’s *t* test (A, C). Data were analyzed using two-way ANOVA, followed by Tukey’s multiple comparisons test (D-M). - c-Jun + CRS versus - c-Jun - CRS *P <0.05, **P <0.01. + c-Jun + CRS versus - c-Jun + CRS, ^#^P <0.05, ^##^P <0.01 (D, I-J).

**Supplementary Figure 7.**
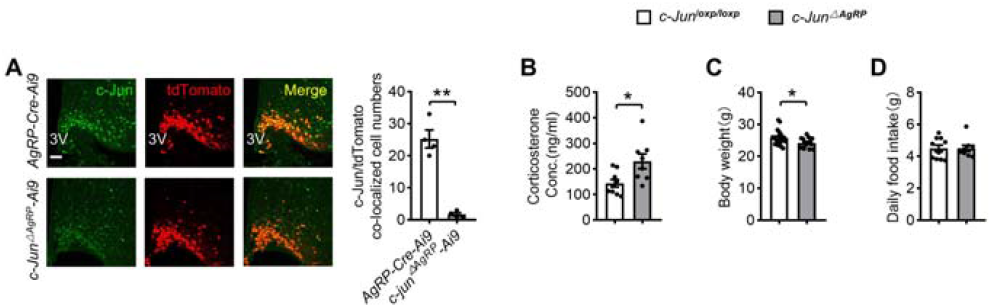
Biochemical parameters related to *c-Jun*^ΔAgRP^ mice. (A) IF staining of tdTomato (red), c-Jun (green) and merge (yellow) in ARC sections (left), and quantification of c-Jun and tdTomato colocalized cell numbers (right). (B) Serum corticosterone levels. (C) Body weight. (D) Food intake. Study for A was conducted using AgRP-Cre-Ai9 mice and *c-Jun*^ΔAgRP^-Ai9 mice. B-D were conducted using c*-Jun*^loxp/loxp^ mice or *c-Jun*^ΔAgRP^ mice. Values are expressed as means ± SEM (n=4-20 per group), with individual data points. Data were analyzed using two-tailed unpaired Student’s *t* test.

**Supplementary Figure 8.**
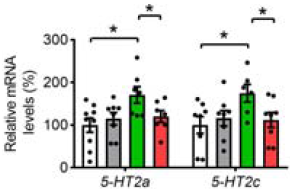
The efficacy of anxiolytic drug cyamemazine (CYA). mRNA levels of 5-HT2a and 5-HT2c. Study was conducted using 18- to 20-week-old *c-Jun*^loxp/loxp^ mice and *c-Jun*^ΔAgRP^ mice treated with (+ CYA) or without (- CYA) followed by DSS administration. Values are expressed as means ± SEM (n=6-9 per group), with individual data points. Data were analyzed using two-way ANOVA, followed by Tukey’s multiple comparisons test.

**Supplementary Figure 9.**
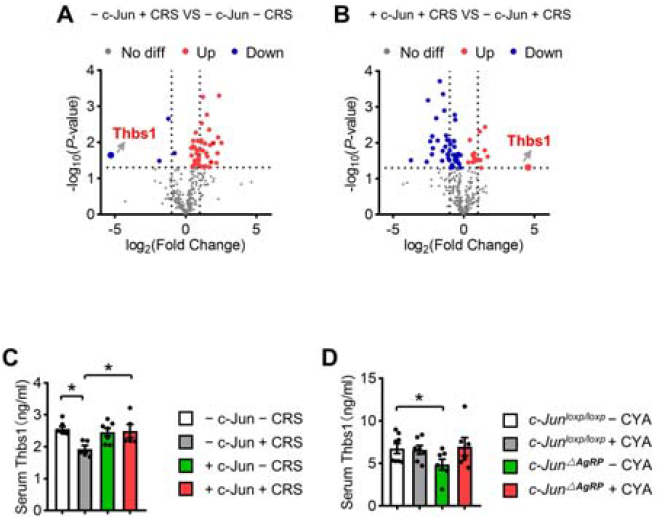
Parameters related to *c-Jun*^ΔAgRP^ mice with thbs1 supplementary. (A) Volcano plot of detected proteins between - c-Jun + CRS mice and - c-Jun - CRS after DSS insults; THBS1 is indicated. (B) Volcano plot of detected proteins between + c-Jun + CRS and - c-Jun + CRS after DSS insults; THBS1 is indicated. (C-D) Serum thbs1 levels. Studies for A-B were conducted using - c-Jun - CRS mice, - c-Jun + CRS mice and + c-Jun + CRS mice to analyze of serum secreted proteins. C was conducted using - c-Jun mice and + c-Jun mice receiving 3% DSS in drinking water for 7 days to induce acute colitis, after stress (+ CRS) or unstress (- CRS). D was conducted using *c-Jun*^loxp/loxp^ and *c-Jun*^ΔAgRP^ mice administrated with 3% DSS for 5 days to induce acute colitis with (+ CYA) or without (- CYA) CYA. Values are expressed as means ± SEM (n=3-7 per group), with individual data points. Data were analyzed using two-tailed unpaired Student’s *t* test (A-B). Data were analyzed using two-way ANOVA, followed by Tukey’s multiple comparisons test (C-D).

